# Idiosyncratic pupil regulation in autistic children

**DOI:** 10.1101/2024.01.10.575072

**Authors:** Isabel Bleimeister, Inbar Avni, Michael Granovetter, Gal Meiri, Michal Ilan, Analya Michaelovski, Idan Menashe, Marlene Behrmann, Ilan Dinstein

## Abstract

Recent neuroimaging and eye tracking studies have suggested that children with autism spectrum disorder (ASD) may exhibit more variable and idiosyncratic brain responses and eye movements than typically developing (TD) children. Here we extended this research for the first time to pupillometry recordings. We successfully completed pupillometry recordings with 103 children (66 with ASD), 4.5-years-old on average, who viewed three 90 second movies, twice. We extracted their pupillary time-course for each movie, capturing their stimulus evoked pupillary responses. We then computed the correlation between the time-course of each child and those of all others in their group. This yielded an average inter-subject correlation value per child, representing how similar their pupillary responses were to all others in their group. ASD participants exhibited significantly weaker inter-subject correlations than TD participants, reliably across all three movies. Differences across groups were largest in responses to a naturalistic movie containing footage of a social interaction between two TD children. This measure enabled classification of ASD and TD children with a sensitivity of 0.82 and specificity of 0.73 when trained and tested on independent datasets. Using the largest ASD pupillometry dataset to date, we demonstrate the utility of a new technique for measuring the idiosyncrasy of pupil regulation, which can be completed even by young children with co-occurring intellectual disability. These findings reveal that a considerable subgroup of ASD children have significantly more unstable, idiosyncratic pupil regulation than TD children, indicative of more variable, weakly regulated, underlying neural activity.

## Introduction

Pupillometry studies in children with ASD have generated considerable interest in recent years (1). Pupil size is regulated by the pupillary light reflex (PLR) (2) and by Norepinephrine (NE) release from the Locus Coeruleus (LC) (3), which also regulates arousal, attention, and exploration/exploitation behaviors (4,5). Hence, potential differences in pupillary responses between ASD and typically developing (TD) individuals could indicate underlying physiological differences in LC-NE and/or PLR function (6).

Several pupillometry studies have reported results from comparisons of relatively small ASD and TD samples (20-30 per group). Studies measuring baseline pupil diameter during stable luminance have reported mixed results with some reporting larger pupil size in ASD relative to TD (7,8) while others have reported the opposite (9), and, yet others report no difference across groups (1,10,11). In contrast, pupil dilation responses to target/novel stimuli with identical luminance (e.g., in oddball tasks) is consistently weaker in ASD (7,8,10), particularly in tasks with higher attentional load (e.g., one-back memory task with added distractors) (11). Smaller pupil dilation suggests weaker LC-NE activity in ASD. Similarly, pupil constriction responses to increased luminance were weaker (i.e., weaker PLR) (12,13) and delayed in time (1) in children and adults with ASD relative to controls. But interestingly, abnormally stronger PLR was reported in 9–10-month-old toddlers who developed ASD at later ages, suggesting different abnormalities during early versus late ASD development (14).

Taken together, the studies above suggest that pupillary responses in ASD children and adults are muted, with LC-NE generating weaker pupil dilations during attention demanding tasks or in response to novel/surprising stimuli, and PLR generating weaker pupil constrictions in response to increased luminance. Because these differential pupillary responses generated by LC-NE (10) and PLR (12,13) are apparent even during passive viewing of sensory stimuli in the absence of a task, they may serve a useful role in distinguishing between ASD and TD individuals in situations where task performance is not possible (e.g., non-verbal children). Hence, there is clear motivation to develop task-free, experimental protocols with child-friendly, engaging stimuli for this purpose.

Previous studies have demonstrated the high ecological validity of using naturalistic movies to study brain function with fMRI (15), especially in young children (16–18). Of particular interest are studies demonstrating that ASD individuals exhibit significantly more variable and idiosyncratic brain responses than TD individuals when observing movies (19,20). These studies quantified idiosyncrasy by measuring inter-subject correlation (inter-SC), which revealed that cortical activity was more strongly correlated across TD individuals observing the same movie than across ASD participants. ASD participants exhibited more idiosyncratic and unique cortical responses with larger between-subject variability. In a recent study we applied the same inter-SC technique to recordings of gaze position during natural viewing of movies. We demonstrated that ASD children also exhibited weaker inter-SC magnitudes, gazing at movies less consistently than TD children and exhibiting more idiosyncratic gaze patterns (21).

Here, we extend this research further by applying the inter-SC approach for the first time to pupillometry data in ASD children. Importantly, we established the largest pupillometry dataset to date from ASD children using an experimental design with 3 different movies, each presented twice, which enabled us to assess the reliability of findings across movies and presentations.

## Methods

A subset of the eye-tracking recordings examined in the current study were analyzed previously to compare gaze patterns across ASD and TD children in a separate study (21). Here, we extracted and analyzed pupillometry data from these and additional recordings.

### Participants

We initially recruited 121 children for the current study through the Azrieli National Centre for Autism and Neurodevelopment Research, between 2016 and 2019. Of these, 81 were diagnosed with ASD according to DSM-V criteria (mean age: 4.46±1.91 years; age range: 1.09-10.07 years; 64 male, 17 female) and 40 were TD (mean age: 4.28±2.10 years; age range: 1.03-10.03 years; 26 male, 14 female). One TD child was removed for having a Social Responsiveness Scale (SRS) score greater than the clinical cut-off (22,23). In addition, data from 15 ASD and an additional 2 TD children were excluded due to low eye tracking quality (see details below), yielding a final sample of 103 children (Table 1). The study was approved by the Soroka Medical Center Helsinki committee and the Ben Gurion University Internal Review Board committee. Written informed consent was obtained from all parents.

**Table 1:** Participant characteristics.

| ***Variable*** | Mean | SD | Range |
| --- | --- | --- | --- |
| **ASD group** (n=66, 53 males) |  |  |  |
| Age at eye tracking (months) | 57.70 | 22.83 | 21-127 |
| ADOS-2 Total CSS (n=52) | 6.69 | 2.65 | 1-10 |
| ADOS-2 SA CSS (n=52) | 6.42 | 2.59 | 1-10 |
| ADOS-2 RRB CSS (n=52) | 7.38 | 2.47 | 1-10 |
| Cognitive Scores (n=46) | 80.39 | 16.80 | 49-117 |
| **TD group** (n=37, 24 males) |  |  |  |
| Age at eye tracking (months) | 53.30 | 22.43 | 15-123 |
| SRS scores (n=34) | 34.21 | 12.68 | 11-57 |

Most of the participating ASD children (52 out of 66) completed the Autism Diagnostic Observation Schedule – 2^nd^ edition (ADOS-2) (24). In addition, 46 of the 66 ASD children completed either a Wechsler Preschool and Primary Scale of Intelligence (25) or the Bayley Cognitive Scales test (26). TD children did not complete ADOS-2 or cognitive tests, but all parents of the TD children, except in three instances, completed the Social Responsiveness Scale (SRS) (22).

### Data acquisition

Participants were seated approximately 60cm from the display screen and the left eye pupil diameter was recorded using an EyeLink 1000+ head-free eye tracking system with a sampling rate of 500 Hz (SR Research Inc. Canada). The participants’ head position was tracked using a sticker placed on their forehead. An infrared camera, located below the display screen, measured pupil size. For each participant, the eye tracker was calibrated prior to data collection: the participant made saccades to each of five stimuli presented sequentially on the screen, and gaze accuracy was then validated to be <2 degrees. Additional validations of calibration accuracy were performed after each movie and re-calibration was performed if error >2 degrees. Experiment Builder and Data Viewer (SR Research Inc. Canada) were used to construct the experiment and visualize the data.

### Experimental design

Once calibration was successfully completed, participants were shown three different movie clips, each presented twice. Each movie was 1.5 minutes long and the total duration of the experiment was roughly 10 minutes. The first movie segment, from the Pixar animation “Jack-Jack Attack”, showed the adventures of a babysitter taking care of an infant with supernatural powers. The second movie segment, taken from the Walt Disney animation “The Jungle Book”, contained a segment in which Mowgli meets the Monkey King who sings and dances while interacting with other monkeys. The third movie contained an un-cut home-video with two sisters (2 and 5 years old) interacting socially in a typical, messy room containing everyday objects.

### Pre-processing and data cleaning

The Eyelink 1000+ records the pupil area as the number of pixels within the image area identified as the pupil, which is equivalent to the angular area of the pupil (27). We identified and removed data segments where the eye tracker lost track of the children’s pupil due to eye blinks and off-screen gazes. This included segments with timepoints where the pupil size equaled zero, was larger than 1200 pixels, or where the pupil size changed faster than 5 pixels per ms – all of which are physiologically implausible. Movies where more than 40% of the data was removed, were entirely excluded from further analysis. We removed 197 of 474 (41.6%) movies observed by ASD children and 43 of 234 (18.4%) movies observed by TD children. This yielded the final sample described above of 66 ASD children and 37 TD children who contributed at least one movie to the analyses.

Of the remaining movies, we removed segments according to the criteria described above such that 16.3±8.6% and 10.9±9.0% of the data were removed from recordings of ASD and TD children, respectively (note that, below, we take the amount of removed data into account in the statistical analysis of the data). We also excluded the first second of each movie to minimize potential stimulus onset responses. Removed time-points were set to NaN values and the remaining segments of analyzed data were smoothed using a Gaussian filter with a width of 250 samples (i.e., half a second). Finally, pupil diameter was computed from angular pupil area using the following formula:

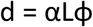

Where d is the pupil diameter, α is a scaling factor deduced empirically by measuring an artificial pupil with known area at a distance of 60cm using our setup (15mm lens, α=0.248), L is the distance between the participant and the screen (i.e., 60cm), and ϕ is the visual angle of the pupil, which is the square root of the angular pupil area measured by the eye tracker (see Eyelink documentation and Hayes and Petrov, 2016).

## Data Analysis

All analyses were performed with custom written code in MATLAB (Mathworks Inc., USA). We extracted time-courses of pupil diameter in mm units for each movie, separately for first and second presentations, per participant. We computed the inter-subject correlation (inter-SC) per movie presentation by correlating the pupil time-courses across all pairs of participants within each group (ASD or TD). We then computed the mean correlation for each child and all other children in their group, yielding a mean inter-SC value per participant. We also computed an intra-subject correlation (intra-SC) value by correlating the pupil time-courses across the two presentations of each movie, per participant.

### Statistical Analyses

To determine whether there were significant differences in the percent of excluded data across groups, we performed a univariate ANOVA analysis with movie type (Jack-Jack Attack, Jungle Book, Naturalistic) as the dependent variable and diagnostic group (ASD and TD) as the between-subjects factor. To evaluate whether there were significant differences in tonic pupil size, variance, inter-SC, and intra-SC across groups, we performed univariate ANCOVAs for each movie with diagnostic group (ASD and TD) as the between-subjects factor, and percent of valid data and age as covariates. Correlations between pupil diameter and behavioral measures (ADOS, cognitive scores) were assessed with Pearson’s correlation coefficient. Finally, we derived receiver operating characteristic (ROC) analyses for the classification of ASD and TD children using the mean inter-SC measure as calculated for each movie per first/second presentation. We computed the Youden index to determine the optimal threshold for separating ASD and TD groups according to the inter-SC values from the first movie presentation and then tested the accuracy of the threshold for separating ASD and TD individuals using the inter-SC values from the second movie presentation.

## Results

Initial analyses demonstrated that there were significantly more excluded/invalid data in recordings of ASD versus TD children in the first presentation of the Jack-Jack Attack (*F*(1)=16.86, *p*=0.0001, η^2^=0.18) and Jungle Book (*F*(1)=10.21, *p*=0.002, η^2^=0.12) movies. We, therefore, included the percent of valid data as a covariate in all further analyses. We also added age as a covariate to ensure that potential differences across groups were not attributable to this variable. The percent of valid data did correlate positively with the age of the ASD children (*r*(64)=0.23, *p*=0.059) but not with their ADOS-2 (ADOS-2 SA: *r*(50)=-0.18, *p*=0.19; ADOS-2 RRB: *r*(50)=-0.23, *p*=0.095; ADOS-2 Total: *r*(50)=-0.19, *p*=0.19) or cognitive (*r*(44)=0.061, *p*=0.69) scores. The ability to contribute valid data in this eye tracking study was, therefore, not significantly associated with the cognitive abilities or core ASD symptom severity of the ASD children.

### No group difference in tonic pupil size

Tonic pupil size, estimated as the mean pupil diameter across all included timepoints of each movie, revealed similar values across ASD and TD participants in all movies and in both presentations (Figure 1). ANCOVA analyses, per movie, demonstrated no significant differences in tonic pupil size across groups for any of the movies (*F*(1)<0.65, *p*>0.42, η^2^<0.009), with no effect of age (*F*(1)<1.22, *p*>0.27, η^2^<0.017) or percent of valid data (*F*(1)<0.86, *p*>0.36, η^2^<0.012). Moreover, the tonic pupil size of individual children in both groups was highly reproducible and significantly correlated across the two presentations of each movie, reflecting the high intra-subject reliability of this measure (TD: *r*>0.93, *p*<0.001; ASD: *r*>0.94, *p*<0.001; Figure 1 B-D).

**Figure 1:**
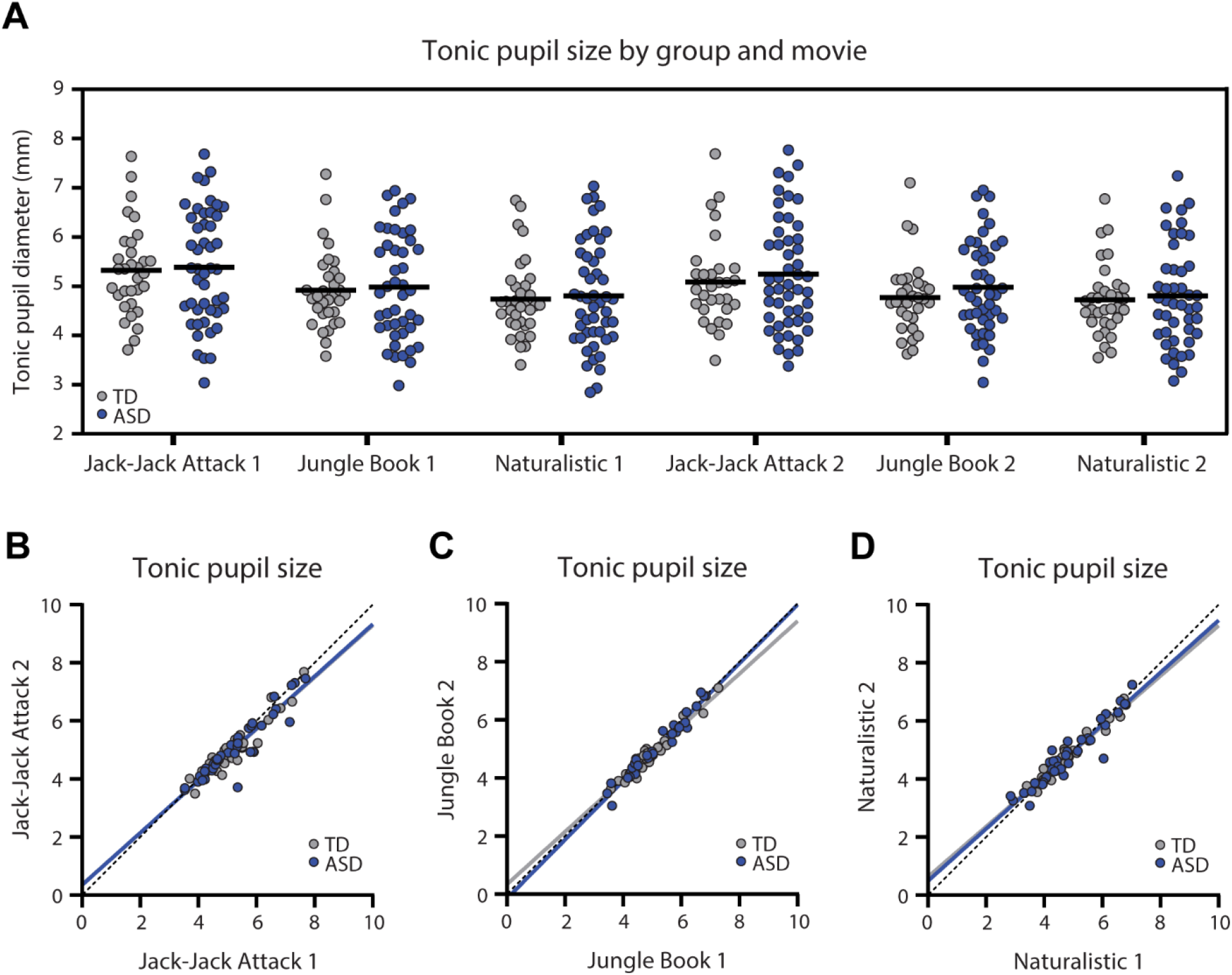
Tonic pupil diameter (mm) in each of the six movies. A. Scatter plot demonstrating tonic pupil diameter for each individual, across both groups, per movie. Black line: mean tonic pupil diameter per group. There were no significant differences across groups. B-D. Scatter plots demonstrating stability of tonic pupil diameter per participant across two presentations of each movie. Each point represents a single child. Solid lines: Least squares fit. Dotted line: unity line. Blue: ASD, Gray: TD.

### Reproducible stimulus-evoked pupillary time-courses

Each movie elicited a unique pupillary time-course, generated by its unique visual content. The mean pupillary time-courses were highly similar across ASD and TD groups per movie (Figure 2) with strong correlations across groups in the Jack-Jack Attack (first presentation: *r*=0.93, *p*<0.001; second presentation: *r*=0.91, *p*<0.001), Jungle Book (first presentation: *r*=0.65, *p*<0.001; second presentation: *r*=0.76, *p*<0.001), and Naturalistic (first presentation: *r*=0.79, *p*<0.001; second presentation: *r*=0.81, *p*<0.001) movies. Correlations were also strong across the two presentations of the Jack-Jack Attack (ASD: *r*=0.89, *p*<0.001; TD: *r*=0.94, *p*<0.001), Jungle Book (ASD: *r*=0.59, *p*<0.001; TD: *r*=0.7, *p*<0.001), and Naturalistic (ASD: *r*=0.51, *p*<0.001; TD: *r*=0.62, *p*<0.001) movies. In contrast, correlations across the different movies were weak and sometimes negative such that, on average, the correlations in both groups were close to zero (TD: *r*=0.03, *p*<0.001; ASD: *r*=0.06, *p*<0.001). While all correlation coefficients were significant due to the large number of degrees of freedom (pupillary time-courses had 44,500 samples), all within-movie correlations were of large effect size (*r*>0.5) while between-movie correlations were of negligible effect size (*r*≤0.06). Taken together, these results demonstrate movie-evoked pupillary time-courses were highly reproducible across presentations and unique to each movie in data from both groups of participants (Figure 2).

**Figure 2:**
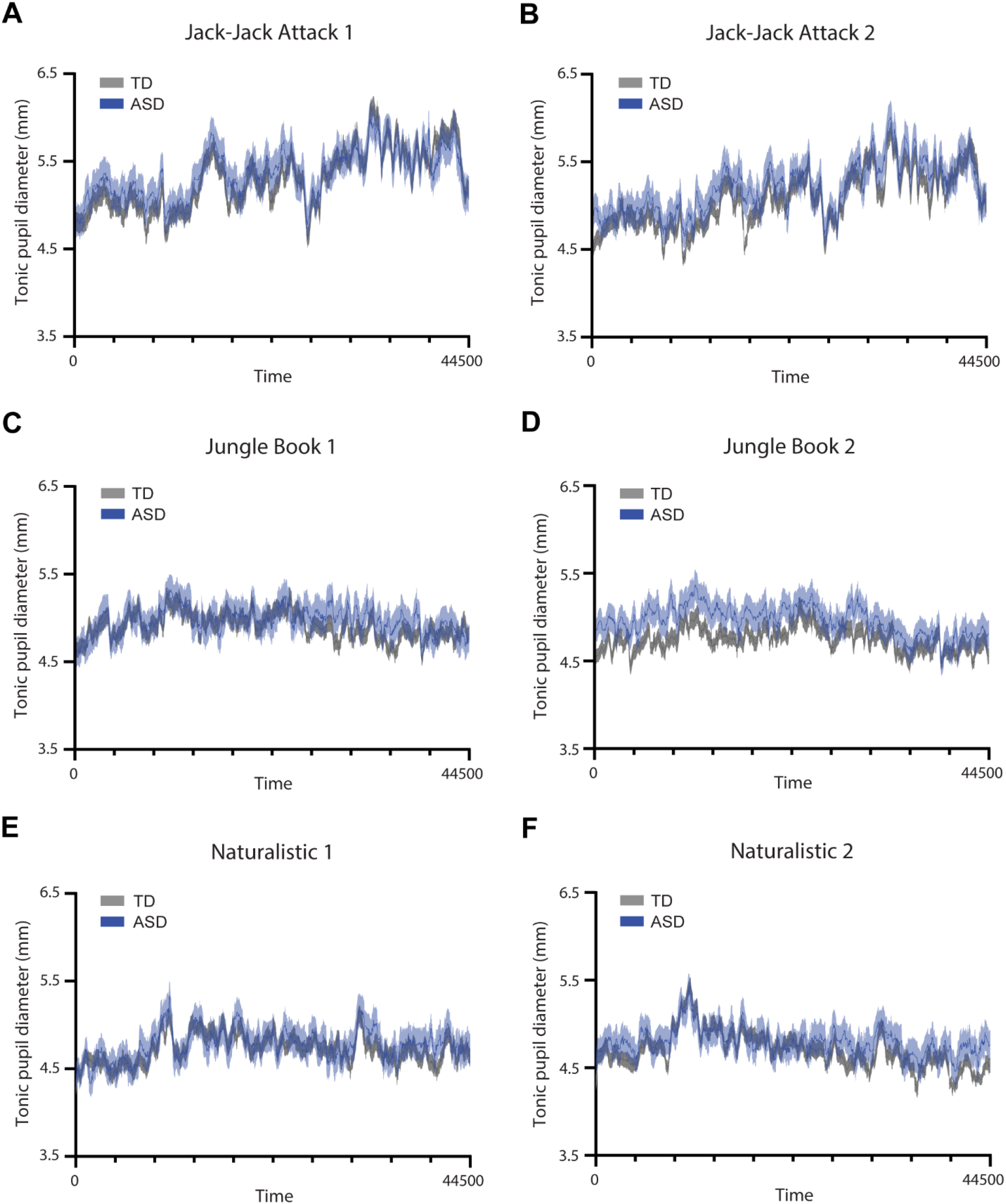
Time-courses of pupil diameter from each presentation of the three movies. A, C, E: First presentation. B, D, F: Second presentation. Solid line: average pupil diameter across participants per group. Shaded area: standard error of the mean. Blue: ASD. Gray: TD.

### Pupillary inter-subject correlations (inter-SC) are consistently weaker in ASD

We computed the pair-wise correlation between the pupillary time-course of a given child and each of the other children in the child’s group, and then computed the mean correlation across all pairs, yielding an inter-SC value per child for each movie presentation (Figure 3). This value represents the similarity of stimulus-evoked pupillary changes between each child and all others in their group.

**Figure 3:**
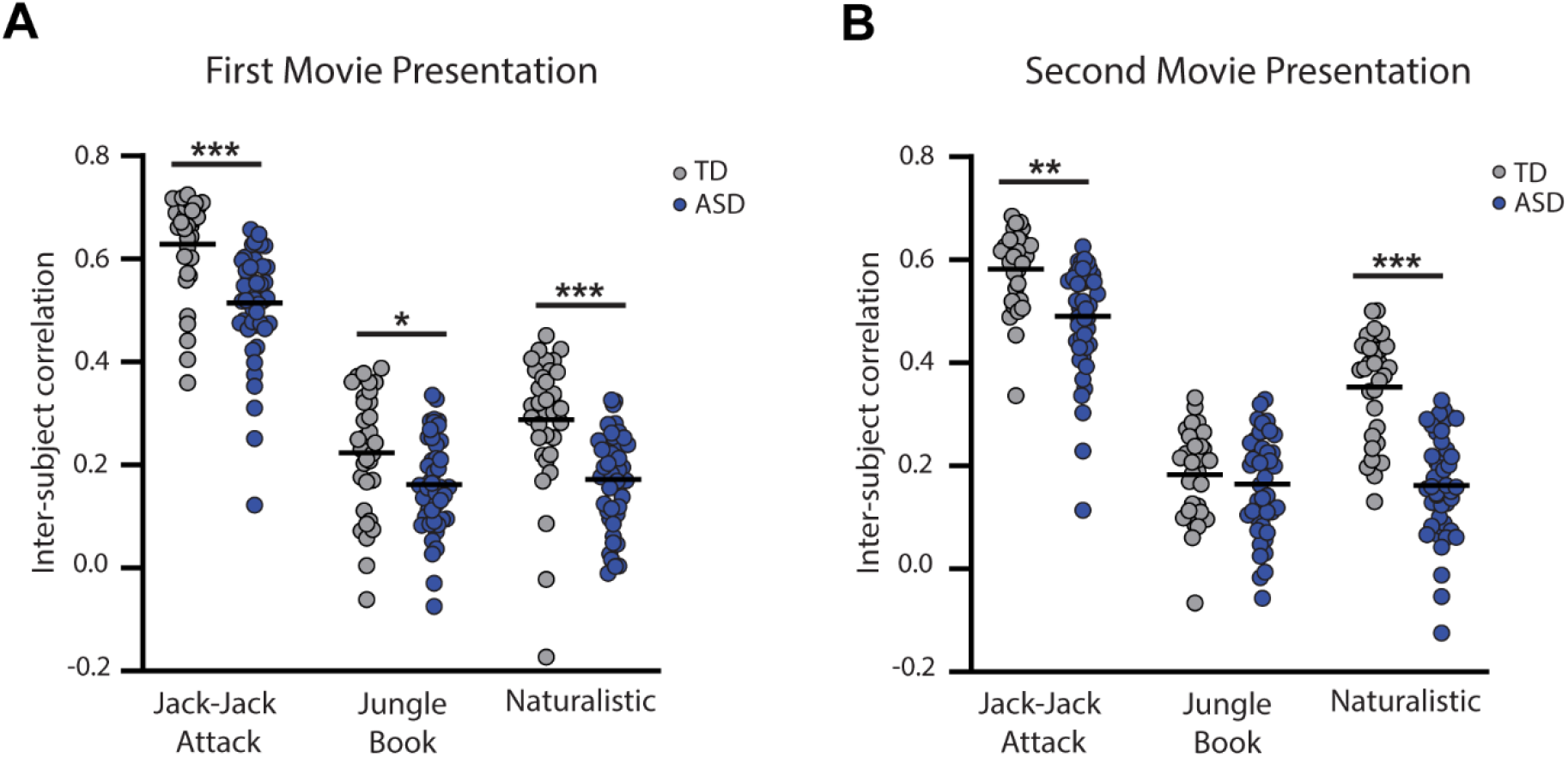
Comparison of inter-subject correlation values across groups for each movie and presentation. A: First presentation. B: Second presentation. Blue: ASD, Gray: TD. Asterisks: Significant differences across groups according to ANCOVA tests (\**p*<0.05, \*\**p*<0.01, *** *p*<0.001).

ANCOVA analyses, with age and percent valid data included as covariates, revealed that inter-SC values were significantly lower in the ASD group in both presentations of Jack-Jack Attack (first presentation: *F*(1)=11.95, p<0.001, η^2^=0.14; second presentation: *F*(1)=9.63, *p*=0.003, η^2^=0.11), both presentations of the Naturalistic movie (first presentation: *F*(1)=16.28, *p*<0.001, η^2^=0.17; second presentation: *F*(1)=56.49, *p*<0.001, η^2^=0.43), and the first presentation of the Jungle Book movie (*F*(1)=6.26, *p*=0.015, η^2^=0.08). There were no significant differences in the second presentation of the Jungle book movie (*F*(1)=0.68, *p*=0.41, η^2^=0.01).

### Pupillary inter-SC are reliable across presentations

Individual inter-SC magnitudes were significantly correlated across presentations of the Jack-Jack (ASD: *r*(34)=0.51, *p*=0.0016; TD: *r*(27)=0.70, *p*<0.001), Jungle Book (ASD: *r*(27)=0.40, *p*=0.032; TD: *r*(25)=0.69, *p*<0.001), and Naturalistic (ASD: *r*(37)=0.62, *p*<0.001; TD: *r*(28)=0.44, *p*=0.015) movie presentations. This demonstrates that the inter-SC measure exhibited relatively high test-retest reliability in both ASD and TD children.

### Classification of ASD and TD children according to their inter-SC

Given the group differences described above, we performed an ROC analysis to quantify the ability of each movie to classify ASD and TD individuals according to their inter-SC values (Figure 5). We compared classification across movies using the Youden Index (*J*) (28), which identifies the inter-SC value (i.e., classification threshold) that yields the highest sum of sensitivity and specificity (optimal point on the ROC). The optimal inter-SC values when analyzing the first presentation of the Jack-Jack Attack, Jungle Book, and Naturalistic movies were 0.64, 0.2, and 0.27, yielding sensitivity/specificity values of 0.96/0.66, 0.66/0.66, and 0.9/0.68, respectively.

Applying these inter-SC thresholds to data from the second movie presentation (i.e., independent sample) yielded sensitivity/specificity values of 1/0.2, 0.52/0.5, and 0.82/0.73, respectively. Taken together, these results suggest that the Naturalistic movie yielded the most reliable between-group classification accuracy.

## Discussion

Our results reveal that ASD children exhibit pupillary responses that are significantly more idiosyncratic (i.e., vary from one individual to another) than those of TD children, consistently across three different movies (Figure 3). The group-average pupillary time-courses were highly correlated across ASD and TD groups (Figure 2) demonstrating that each movie elicited a unique and reliable stimulus-driven pupillary time-course, on average. However, pupillary time-courses of ASD children diverged from the mean (i.e., weaker inter-SC) to a larger extent than those of TD children. Individual magnitudes of inter-SC were strongly correlated across presentations (Figure 4) demonstrating that idiosyncrasy (how different one is from the group) is a reproducible individual characteristic.

**Figure 4:**
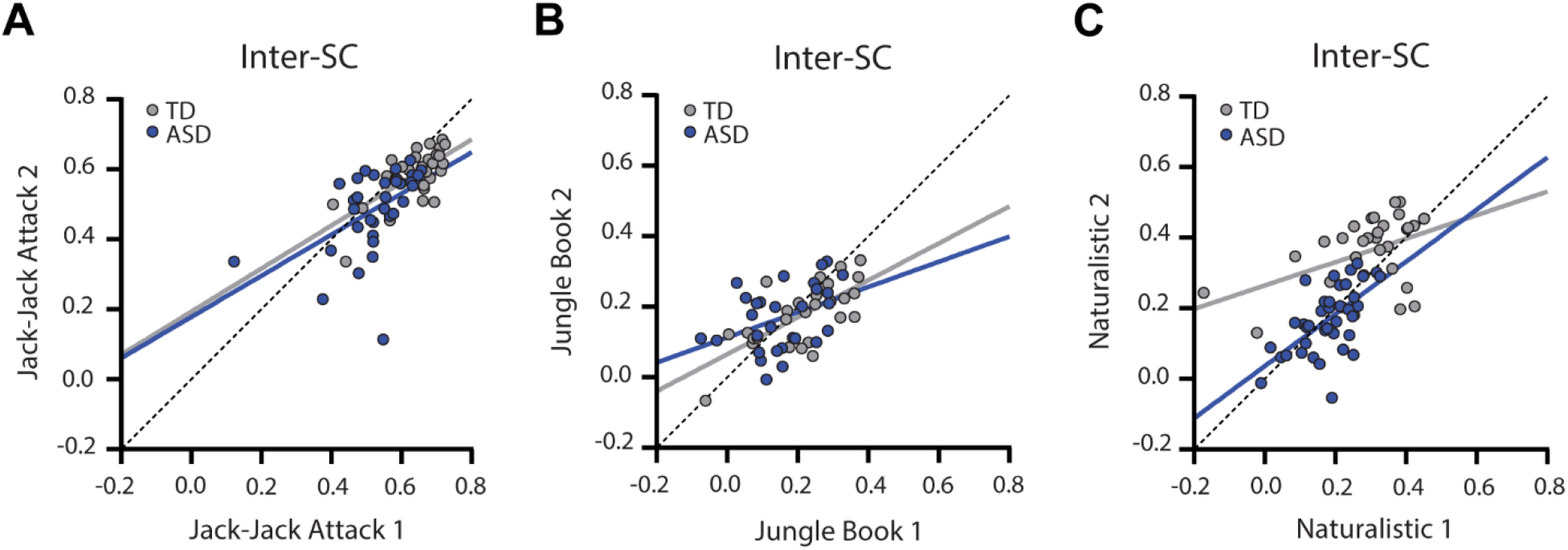
Scatter plots of inter-SC values demonstrating their correlation across the two presentations of Jack-Jack Attack (A), Jungle Book (B), and Naturalistic (C) movies. Blue: ASD, Gray: TD. Solid lines: Least squares fit. Dotted line: unity line.

**Figure 5:**
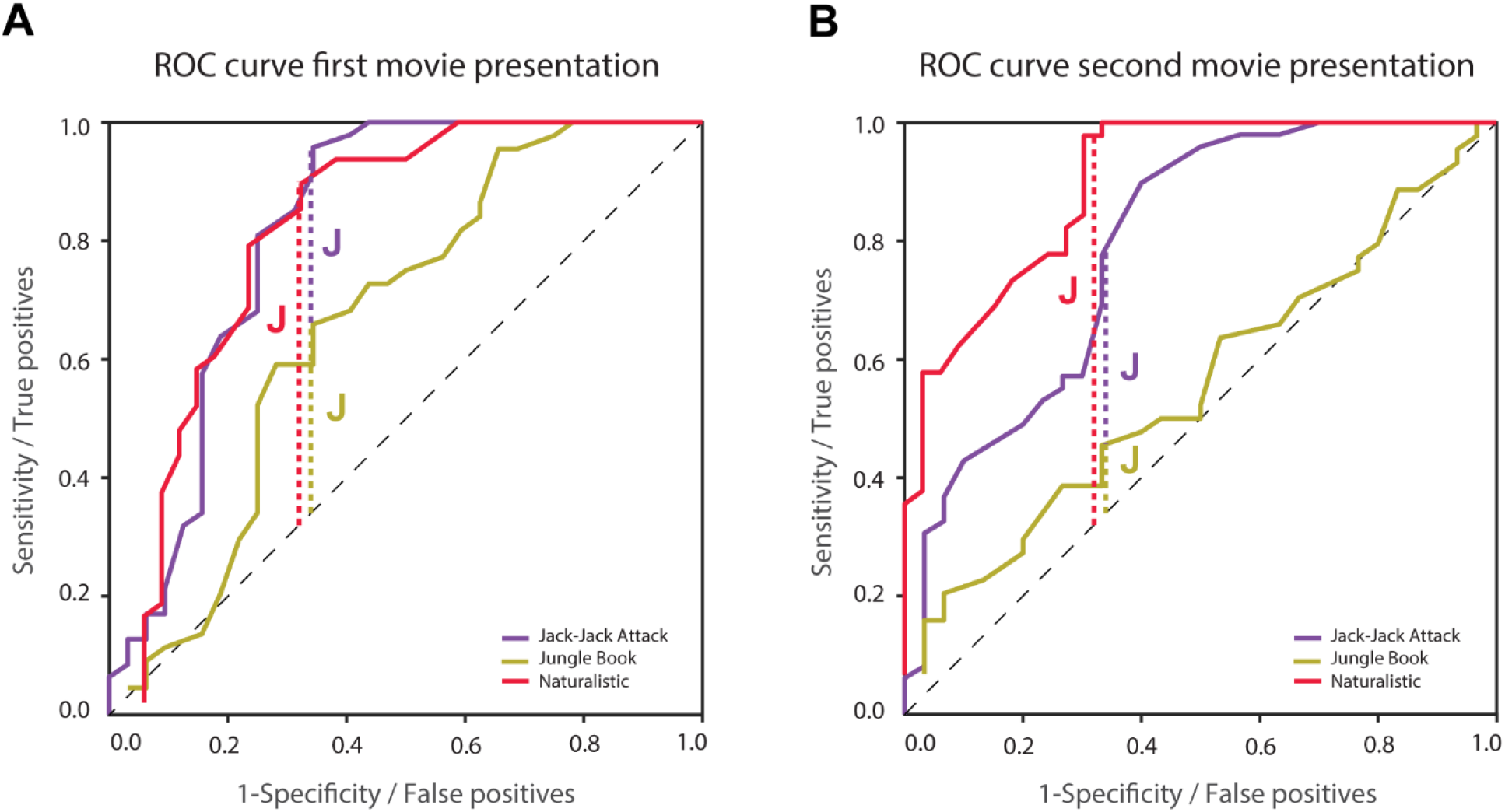
ROC analyses using inter-SC to classify ASD and TD participants. A: First presentation. B: Second presentation. Vertical dashed lines represent the optimal classification threshold as determined using Youden’s J statistic for the Jack-Jack Attack (purple), Jungle Book (yellow), and Naturalistic (red) movies. Optimal classification thresholds were determined with data from the first presentation (left panel) and their accuracy was tested with data from the second presentation (right panel). Black dashed line: unity line.

Inter-SC differences were most pronounced in the pupillary responses to a naturalistic un-cut home-video of two TD children engaged in a social interaction. Extracting a simple inter-SC threshold based on ROC analysis and Youden Index using data from the first presentation of this movie, enabled us to classify ASD and TD children with 82% sensitivity and 73% specificity in held-out independent data from the second movie presentation (Figure 5). These results highlight the potential clinical utility of the inter-SC measure for identifying many ASD individuals when using pupillometry recordings during movies with naturalistic social interactions.

These results indicate that pupil size regulation by LPR and LC-NE mechanisms is more variable (i.e., less reliable) in children with ASD than in TD controls. Such an interpretation would be consistent with previous hypotheses proposing that sensory neural responses in the visual, auditory, and tactile domains may be more variable in some ASD individuals (29,30).

### Idiosyncrasy rather than consistent group differences

A growing body of literature demonstrates that ASD individuals exhibit idiosyncratic behavioral and physiological responses that differ from TD responses. Examples include idiosyncratic fMRI activation time-courses in response to movies (19,20), idiosyncratic resting-state fMRI functional connectivity (31–33), idiosyncratic gaze patterns when observing movies (21,34), as well as other behavioral idiosyncrasies (35,36). These studies suggest that rather than exhibiting consistently weaker or stronger behavioral or physiological responses, ASD individuals tend to differ from one another to larger extents and in unique ways. This means that, on average, measures may be similar to those of TD individuals, yet ASD individuals are scattered further away from the mean.

Quantifying individual idiosyncrasy, as calculated here with inter-SC, reveals that the majority, but not all ASD participants exhibit large idiosyncrasy. Our results demonstrated that idiosyncratic pupillometry responses to the Naturalistic movie (defined by a classification threshold of inter-SC<0.27) were apparent in 82% of the ASD participants and only 27% of the TD participants (see ROC analysis). For comparison, a previous fMRI study reported that about one third of ASD participants exhibited distinctive idiosyncratic brain activations when observing a movie (20). In the context of the current study, the high selectivity and specificity for classifying ASD/TD group membership suggests that quantification of pupillometry idiosyncrasy may be useful for identifying a large group of ASD children with poorer LPR and LC-NE regulation.

Importantly, the approach outlined here does not require cooperation or compliance with an explicit task, which may be difficult for the more severe or younger ASD individuals, thereby extending its utility and generalizability to the broader ASD population. Future studies could assess whether idiosyncrasy magnitudes are similarly apparent across multiple behavioral and physiological domains and multiple sensory modalities. This could reveal potential generalized behavioral and neural idiosyncrasy or specific idiosyncrasy in particular domains, in distinct subgroups.

### Pupillometry abnormalities in ASD

A variety of studies have reported that ASD participants may exhibit different pupillometry abnormalities indicative of underlying hyper- or hyporegulation by LC-NE and/or LPR circuits. One hypothesis is that ASD participants may exhibit larger tonic pupil size indicative of stronger LC-NE tonic activity, potentially associated with hyper-arousal and increased behavioral flexibility (5). While some studies with relatively small samples (<32 ASD participants in each) have reported significantly larger tonic pupil size in ASD (7,8), others have reported the opposite (9) or no differences across groups (1,10,11). Our results, with a somewhat larger sample and with multiple, repeated measurements, suggest that there is indeed no significant difference in baseline pupil size across groups (Figure 1).

Another hypothesis is that ASD individuals may exhibit weaker pupil dilations to stimuli with cognitive, attentional, or social load that may indicate poor phasic LC-NE modulation. Phasic increases in LC-NE dopamine innervation are important for increasing arousal during task demanding periods and achieving optimal performance (5). Several studies have reported that pupillary responses in ASD individuals are weaker to visual stimuli in the context of a spatial attention task (8), an odd-ball task (7), a one-back memory task (11), or when stimuli include social information (37,38).

A third hypothesis suggests that ASD individuals may exhibit weaker pupil constriction in response to luminance increases (i.e., weaker PLR). Several studies have indeed reported weaker (12,13) and delayed (1) PLR responses in children and adults with ASD. Surprisingly, there have also been reports of abnormally strong PLR responses in 9–10-month-old toddlers who developed ASD at later ages(14).

Taken together, the last two hypotheses suggest attenuated pupillometry time-courses in ASD with weaker PLR associated constrictions and weaker LC-NE associated dilations. Our pupillometry results do not seem to show differences, on average, in the pupillary response time-courses across ASD and NT groups, in any of the movies (Figure 2). Pupillary time-courses were highly correlated across groups, demonstrating that the temporal structure of pupillary responses was overall highly similar across groups. Nevertheless, our study was not designed to separate PLR and LC-NE responses or isolate specific stimulus events that would be expected to generate a pupillary response. Hence, our results do not offer strong evidence for the existence of specific LC-NE and LPR differences across groups or lack thereof. Further studies are necessary to test hypotheses regarding potential atypicalities in each of these neural mechanisms in ASD.

### Limitations

While our study utilized a highly ecological design that enabled inclusion of ASD children across the spectrum including those with intellectual disability, it had several important limitations. First, we did not design our stimuli or analyze it post-hoc to separate pupillary responses associated with PLR versus LC-NE mechanisms. It is likely that naturalistic, complex stimuli contain multiple transitions in both low-level visual features (e.g., luminance and contrast) as well as high-level content features (e.g., narrative complexity, novelty, social and emotional valence). Hence, our study measured pupillary changes that were the product of both PLR and LC-NE regulation. Second, data loss was clearly an issue in the current study. While all 103 children included in the study successfully watched at least one movie, most participating children did not successfully watch all 6 movies. Since our experimental design included considerable redundancy with multiple movies and presentations, we believe that our results are reliable and conclusive despite partial data collection from many children. Nevertheless, this raises an important limitation of eye tracking studies with ASD children. While previous studies have reported success rates as high as 95% in collecting eye tracking data from ASD children (39), these were collected from ASD children without intellectual disability. We believe that improving stimuli and acquisition conditions to maximize data collection from young children with ASD and intellectual disability should be an important goal of future studies.

### Conclusions

Most children with ASD exhibit pupillometry time-courses that differ from those of TD children. This suggests that pupil regulation differs in ASD children in idiosyncratic ways, which can be quantified using inter-SC, including in children with ASD and intellectual disability. Beyond the clinical value of this measure for identifying individuals with ASD, it suggests that PLR and LC-NE circuits do not operate in a uniform fashion across ASD individuals. Rather than attenuated or excessive circuit responses, we speculate that these circuits may exhibit larger variability in ASD individuals, perhaps due to excitation/inhibition imbalances that have been implicated as a potential underlying mechanism (40,41). Further studies delineating PLR and LC-NE responses while using naturalistic stimuli that can be used with large samples of ASD children, including those with intellectual disability, are highly warranted.

## Acknowledgements

This research was supported by Fulbright U.S. Student Program to IB, Israeli Science Foundation (1150/20), Israeli Ministry of Science and Technology, and Azrieli Foundation grants to ID, National Institute of General Medical Sciences grants T32GM008208 to IB and MCG, T32GM081760 to MCG, and a Simons Foundation Autism Research Initiative grant to MB. MB acknowledges support from P30 CORE award EY08098 from the National Eye Institute, NIH, and unrestricted supporting funds from The Research to Prevent Blindness Inc, NY, and the Eye & Ear Foundation of Pittsburgh. The content is solely the responsibility of the authors and does not necessarily represent the official view of the Fulbright Program, the National Institute of General Medical Sciences, or the National Institutes of Health. Finally, the authors thank the participants and their families for making this research possible.

